# Profile of Molecular Subtypes of Breast Cancer Among Bangladeshi Women - Audit of Initial Experience

**DOI:** 10.1101/253781

**Authors:** Md. Zillur Rahman, Anwarul Karim

**Affiliations:** Head of the Department and Associate Professor, Department of Pathology, Chittagong Medical College, Chittagong, Bangladesh; Intern doctor, Chittagong Medical College and Hospital, Chittagong, Bangladesh

**Keywords:** Breast cancer, molecular subtypes, immunohistochemistry, HER2, luminal A, luminal B, basal-like

## Abstract

**Background:** Receptor status and molecular subtyping of breast cancer are crucial for patient management. We present here our initial experience on the status of different molecular subtypes and clinicopathological characteristics of invasive breast carcinomas in Bangladeshi population especially in Chittagong zone.

**Materials and methods:** A total of 59 histopathologically confirmed cases of invasive ductal carcinoma were selected for this study. Fifteen out of 59 cases were reported as HER2 equivalent and could not be categorized into any subtype because of the lack of availability of fluorescence in situ hybridization. The remaining 44 cases were distributed into different molecular subtypes and then the clinicopathological characteristics were compared for each molecular subtype.

**Results:** Age ranges from 24-70 years with a mean age of 43.95 years. Most of the patients were in 41-50 years age group. Among the 44 cases, most common subtype was HER2/neu amplification (13 cases, 29.55%). Luminal A, luminal B and basal like subtypes were 11 (25%), 10 (22.73%) and 10 (22.73%) respectively. The mean tumor size was 3.46 cm and the highest mean tumor size was in basal-like subtype (4.01cm). Twenty five out of 59 cases (42.37%) showed axillary lymph node metastasis. Lowest axillary lymph node metastasis was found in luminal A subtype (3/11=27.27%).

**Conclusion:** HER2/neu amplification subtype was found to be more common in this region. Luminal A subtype was found to be more favorable in comparison to the other subtypes in terms of axillary lymph node metastasis.

## INTRODUCTION

Carcinoma of breast is the most common non-skin malignancy in women and is second only to lung cancer as a cause of cancer deaths [1]. Variations in gene expression patterns in breast tumors have been identified by using complementary DNA microarray technology which defines several different molecular subtypes [2]. DNA microarray technology is very expensive and cannot be used on formalin-fixed, paraffin-embedded samples. Recent findings indicate that immunohistochemical (IHC) protein expression profiles are surrogates for intrinsic gene derived expression profiles defining molecular subtypes of breast cancer [3]. Molecular subtyping is crucial because these subtypes show striking differences with regard to patient characteristics, clinicopathological features, treatment response, and outcome [1]. In this study, we grouped our cases based on IHC markers into four subtypes - a) Luminal A: Estrogen Receptor (ER) and/or Progesterone Receptor (PR) positive and Human Epidermal Growth Factor Receptor 2 (HER2) negative, b) Luminal B: ER and/or PR positive and HER2 positive, c) HER2/neu amplification: ER and PR negative and HER2 positive and d) Basal like: ER, PR and HER2 all negative [Table-1]. This classification is an adaptation of original Carey method, where CK5/6 and HER1 were also incorporated in addition to ER, PR, and, HER2 [4]. Luminal B, HER2/neu amplification and basal like subtypes are known to be more clinically aggressive and have poorer prognoses compared with luminal A tumors [5,6,7]. The prevalence of molecular subtypes of breast carcinomas and their clinicopathological characteristics have not been studied extensively in Bangladesh yet. The prime objective of this study was to estimate the status of different molecular subtypes of invasive breast carcinomas in Bangladeshi population especially in Chittagong zone. In addition, we also compared some clinicopathological characteristics i.e. age, tumor size and axillary lymph node status of each molecular subtype.

**Table 1:** Different molecular subtypes based on IHC ER, PR and HER2 status

| **Molecular subtypes** | **Immunohistochemical characterization** |
| --- | --- |
| Luminal A | ER and/or PR positive, HER2 negative |
| Luminal B | ER and/or PR positive, HER2 positive |
| HER2/neu amplification | ER negative, PR negative, HER2 positive |
| Basal like | ER negative, PR negative, HER2 negative |

## MATERIALS AND METHODS

A total of 59 histopathologically confirmed cases of breast cancer were selected. These cases were diagnosed in the department of pathology of Chittagong medical college and hospital (CMCH) and in the Care investigation histopathology laboratory, Chittagong from resected breast with or without axillary lymph nodes referred from different hospitals over a period of seven months (January 2015 – July 2015). All the received formalin fixed specimens were examined grossly and the representative samples were processed for paraffin sections; and then stained with hematoxylin and eosin (H & E). IHC was done using prediluted ready to use antibodies. For ER and PR, Dako monoclonal mouse anti-human ER, clone 1D5 and Dako monoclonal mouse anti-human PR, clone PgR 636 were used respectively and reporting was performed according to Allred guidelines [8]. For HER2, Dako rabbit antihuman HER2 protein was used and reporting was performed according to ASCO/CAP guideline (2013) [9]. All the cases were of invasive ductal carcinoma. We found that 15 out of 59 cases were HER2 equivocal (score 2+). These IHC HER2 equivocal cases could not be further investigated for gene amplification by fluorescence in situ hybridization (FISH), as this expensive test is not available in this region and as a result we could not categorize them into any molecular subtype. We distributed the remaining 44 cases into four molecular subtypes based on IHC ER, PR and HER2 status as shown in Table-1. We assessed age, tumor size and axillary lymph node metastasis for each molecular subtype.

## RESULTS

The patients were from 24-70 years of age with a mean age of 43.95 years. Forty-one to fifty years age group represented most of the patients (25/59=42.37%) followed by 31-40 years (17/59=28.81%), 51-60 years (7/59=11.86%), 21-30 years (6/59=10.17%) and 61-70 years age groups (4/59=6.78%) [Table-2].

Among the molecular subtypes, HER2/neu amplification subtype was the most common (13/44=29.55%) [Figure-1]. Luminal A subtype was found mostly in 31-40 years age group (5/11=45.45%). Luminal B had 30% (3/10) cases in 31-40 years age group and also 30% (3/10) cases in 41-50 years age group. HER2/neu amplification and basal like subtypes were found mostly in 41-50 years age groups, 46.15% (6/13) and 60% (6/10) respectively.

**Figure 1:**
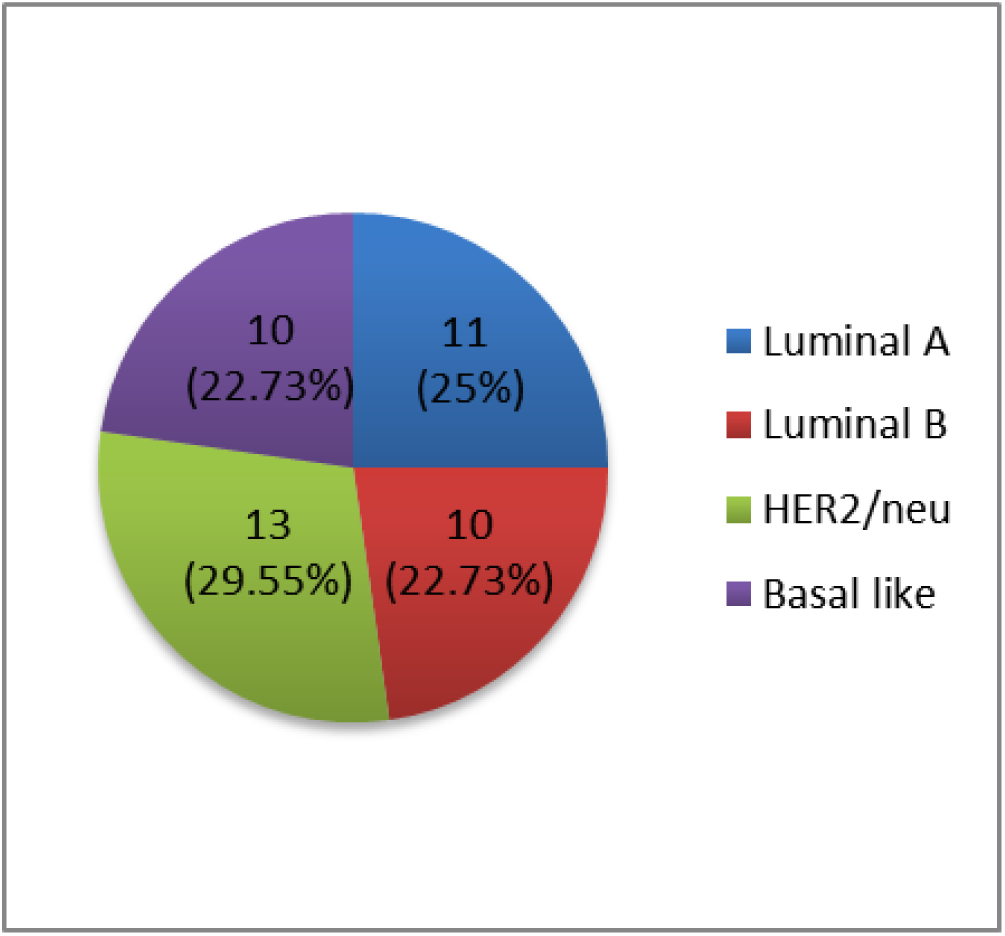
Distribution of cases according to different molecular subtypes

The mean tumor size was 3.46 cm ranging from 1.4 cm to 9 cm in their largest dimension on gross examination. 10.17% tumors (6/59) were of less than 2 cm, 81.36% tumors (48/59) were in the range of 2-5 cm and 8.47% tumors (5/59) were more than 5 cm. All the tumors in basal like subtype (10/10) were more than 2 cm in size followed by luminal A subtype with 90.91% (10/11). Luminal B and HER2/neu amplification subtypes had 80% (8/10) and 76.92% (10/13) cases which were more than 2 cm in size. Among the molecular subtypes basal like was found to have the highest mean tumor size (4.01 cm) followed by HER2/neu amplification (3.67 cm), luminal A (3.05 cm) and luminal B subtypes (2.99 cm) [Table-2].

**Table 2:** Clinicopathological characteristics of breast tumors according to molecular subtypes

| Clinicopathological characteristics |  | Luminal A<br>(ER and/or PR+, HER2-)<br>(11/59) | Luminal B<br>(ER and/or PR+, HER2+)<br>(10/59) | HER2/neu amplification<br>(ER-, PR- and HER2+)<br>(13/59) | Basal like<br>(ER-, PR-, HER2-)<br>(10/59) | HER2 equivocal (score 2+) |  | Total<br>(59) |
| --- | --- | --- | --- | --- | --- | --- | --- | --- |
|  |  |  |  |  |  | ER and/or PR+<br>(4/59) | ER and PR-<br>(11/59) |  |
| Age (years) | 21-30 | 1 (9.09%) | 1 (10%) | 1 (7.69%) | 1 (10%) | 1 | 1 | 6 (10.17%) |
|  | 31-40 | 5 (45.45%) | 3 (30%) | 4 (30.77%) | 1 (10%) | 0 | 4 | 17 (28.81%) |
|  | 41-50 | 3 (27.27%) | 3 (30%) | 6 (46.15%) | 6 (60%) | 3 | 4 | 25 (42.37%) |
|  | 51-60 | 2 (18.18%) | 2 (20%) | 1 (7.69%) | 2 (20%) | 0 | 0 | 7 (11.86%) |
|  | 61-70 | 0 | 1 (10%) | 1 (7.69%) | 0 | 0 | 2 | 4 (6.78%) |
|  | Total | 11 (100%) | 10 (100%) | 13 (100%) | 10 (100%) | 4 | 11 | 59 (100%) |
| Tumor size (cm) | <2 | 1 (9.09%) | 2 (20%) | 3 (23.08%) | 0 | 0 | 0 | 6 (10.17%) |
|  | 2-5 | 10 (90.91%) | 7 (70%) | 9 (69.23%) | 9 (90%) | 4 | 9 | 48 (81.36%) |
|  | >5 | 0 | 1 (10%) | 1 (7.69%) | 1 (10%) | 0 | 2 | 5 (8.47%) |
|  | Total | 11 (100%) | 10 (100%) | 13 (100%) | 10 (100%) | 4 | 11 | 59 (100%) |
| Mean tumor size (cm) |  | 3.05 | 2.99 | 3.67 | 4.01 | 3.08 | 4.02 | 3.4 |
| Axillary lymph node metastasis | 0 | 8 (72.73%) | 6 (60%) | 8 (61.54%) | 6 (60%) | 1 | 5 | 34 (57.63%) |
|  | <4 | 3 (27.27%) | 3 (30%) | 3 (23.08%) | 3 (30%) | 1 | 5 | 18 (30.51%) |
|  | ≥4 | 0 | 1 (10%) | 2 (15.38%) | 1 (10%) | 2 | 1 | 7 (11.86%) |
|  | Total | 11 (100%) | 10 (100%) | 13 (100%) | 10 (100%) | 4 | 11 | 59 (100%) |

Twenty five out of 59 cases (42.37%) showed axillary lymph node metastasis. Axillary lymph node metastasis in luminal A subtype was 27.27% (3/11). This was significantly lower than that of luminal B (40%=4/10), HER2/neu amplification (38.46%=5/13) and basal like subtypes (40%=4/10%) [Table-2].

## DISCUSSION

The study population in our study ranged from 24-70 years with a mean age of 43.95 years. Bennis et al. [10] reported mean age of 45 years in a similar study of 366 cases in Morocco. Akbar et al. [11] found mean age of 47.55 years in a study of 60 cases in Pakistan. The mean age of our study population nearly corresponds with that of the study population of Bennis et al. [10] and Akbar et al. [11]. However, Zhu et al. [12] found a mean age of 51 years in a similar study of 3198 cases in China. The lower mean age in our study population, Morocco and Pakistan than China can be explained by lower life expectancy in these countries compared with China. Moreover, Zhu et al. [12] worked on a larger sample size compared to others.

In this study, HER2/neu amplification subtype was the most common subtype comprising 29.55% (13/44) of the total cases. Akbar et al. [11] found this subtype to be the most prevalent in their study in Pakistan (30%) as well. A study of 112 patients in India by Kumar et al. [13] found that HER2/neu amplification subtype was present in 46.37% cases. It indicates that HER2/neu amplification subtype in Bangladesh, Pakistan and India is significantly higher than what most western literatures report e.g. in the study by Hawladar et al. [14] (only 4.6%). On the other hand, the study by Hawladar et al. [14] found luminal A subtype to be of highest incidence (72.7%). Bennis et al. [10] and Zhu et al. [12] also reported luminal A subtype to be the most prevalent, 53.6% and 65.3% respectively. But in our study, luminal A subtype showed significantly lower incidence which is only 25% (11/44). This lower incidence of luminal A subtype in the present study may be partly due to less frequent use of menopausal hormone therapy and lack of mammographic screening program. But this demands further study to establish these factors as causes of lower incidence of luminal A subtype. Another possible reason may be that, since it is not a population based study rather a study on some selected case series, we may not get the exact scenario.

The mean tumor size in our study was 3.46 cm and 89.83% tumors were of more than 2 cm size in their largest dimension. Kumar et al. [13] from India also found similar results. They reported mean tumor size 3.4 cm and 85.8% of their cases were having tumor size more than 2 cm. However, Zhu et al. [12] reported mean size of 2.1 cm. The higher mean tumor size in our study and in India may be due to late consultation during the progression of the disease because of the existing social circumstances in this subcontinent. Another important cause may be the lack of mammographic screening program. All the basal like tumors (10/10) in this study were of more than 2 cm in size and the mean size of tumors was also highest in this subgroup (4.01 cm). This result corresponds with the result of Kumar et al. [13] where all the basal like tumors were of more than 2 cm size as well.

Our study found lowest axillary lymph node metastasis rate in luminal A subtype (27.27%=3/11). Luminal B, HER2/neu amplification and basal like subtypes showed nearly equal lymph node metastasis rate. Kumar et al. [13] and El-Hawary et al. [15] in Egypt also found luminal A subtype to have the lowest rate of lymph node metastasis.

We could not categorize 15 out of 59 cases to any specific molecular subtype because of their IHC HER2 equivocal status. Reflex testing using FISH should have been performed for those HER2 equivocal cases according to the recommendation by ASCO/CAP guideline (2013) [9] which would have given more precise results of prevalence of molecular subtypes in this area. Unfortunately, FISH is not available here. Although very recently FISH has been instituted in a center in Bangladesh but it is still very expensive and inadequate to serve the large Bangladeshi population. Hence, FISH should be made more available that will guide the oncologists to decide whether to advice HER2 targeted therapy or not for immunohistochemically HER2 equivocal cases as some of these may show gene amplification in FISH.

## CONCLUSION

Our study found that 4th & 5th decades are the most affected age groups by breast carcinoma in this region. The mean size of the tumors and axillary lymph node involvement were found to be high in this study. HER2/neu amplification subtype was found to be more common in this region than in the western countries. Luminal A subtype was shown to be more favorable in comparison to the other subtypes in terms of axillary lymph node metastasis. FISH should be made available in this region as it guides treatment options, especially in HER2 equivocal cases.

## LIMITATION

This paper was prepared from the data of our initial observations. It lacks sufficient statistical power to draw any definite conclusion due to the limited sample size. Nevertheless, this study provides some insight about different molecular subtypes and clinicopathological features of breast cancer in Chittagong.

